# Isolation and identification of *Phytophthora* species оn nut-bearing plants

**DOI:** 10.1101/2020.12.21.423714

**Authors:** T. A. Surina, M. B. Kopina, A.V. Smirnova, D.I. Shukhin

**Affiliations:** All-Russian plant quarantine center (FGBU “VNIIKR”)

## Abstract

The paper describes the isolation and identification of pathogens from soil samples taken in the Moscow region from walnut plants with signs of wilting and stem ulcers. Isolation of *Phytophthora* from soil samples was carried out by the method of floating biological baits with subsequent isolation of pathogens on a semi-selective nutrient medium - P5ARP [H]. Samples were taken from the colonies for DNA isolation. After that, PCR analysis was carried out with primers YPh1F / YPh2R and ITS4 / ITS5 and sequencing. As a result of the studies carried out, colonies similar to the morphological characteristics of *P. cactorum* and *P citricola* were obtained and confirmed by sequencing.

## Introduction

The genus *Phytophthora* is one of the largest in the order *Peronosporales*, includes more than 180 described species, which differ in the degree of pathogenicity and harmfulness both in agricultural and ecological terms. Over the past few decades, interest in oomycete data has noticeably increased, including the introduction of molecular genetic studies reflected in the increased number of described taxonomic units: from 58 species in 1996 to more than 150 species in 2017 (Yang X., 2017). The causative agents of late blight are soil fungus-like organisms from the genus *Phytophthora* infect a wide range of host plants, causing foot rot and root rot, leaf burn, shoot death, bark necrosis, fruit rot, ulcerative lesions and other manifestation symptoms.

*Phytophtora* remains viable in soil for a long time. Pathogens are spread by zoospores by water flows during irrigation or as a result of heavy rains; they can also be spread by soil particles containing oospores, which are carried on agricultural implements or machines.

Diagnosis of *Phytophthora* species is fundamental to control and prevent the spread of these pathogens (Cooke et al., 2007). Direct isolation of *Phytophthora* spp. from the soil is hindered by the biological features of these pathogens, they have a rather slow growth and are easily inhibited by accompanying micromycetes.

Selective media for the isolation of pathogens usually consists of corn agar (CMA) or V8 agar with different combinations of antibiotics: pimaricin, ampicillin, rifampicin, vancomycin, nystatin; and fungicides: pentachloronitrobenzene (PCNB) and hymexazole. The selective action of these agents is mainly provided by pimaricin or nystatin, since they are active against most fungi from the *Eumycota* group. Selective media are light sensitive and should be stored in the dark. Gimexazole inhibits most of the *Pythium* and *Morteriella* species, which can inhibit and mask *Phytophthora*. The combination of rifampicin and ampicillin is more effective in suppressing bacteria than vancomycin.

A wide range of methods have been reported for the successful isolation of *Phytophthora* from soil and water samples, including direct seeding of soil onto the medium and the bait method. Due to the limited amount of soil that can be applied to the surface of the medium, direct seeding is usually used when the soil is heavily infested with *Phytophthora*. The bait method has several advantages over direct seeding. Firstly, a larger volume of soil can be tested, which increases the likelihood of detecting species present in the sample. Secondly, homothallic species that persist as dormant oospores are more likely to be detected by bait techniques than by direct seeding.

Currently, there are reliable molecular research methods that make it possible to identify the pathogen at any stage of its development, and the availability of data on regions of the genome allows the development of new modern methods of identification.

## Materials and methods

The selection of soil samples was carried out in the Michurinsky orchard of the Russian State Agrarian University named after K.A. Timiryazev. In total, 29 soil samples were taken from plants of walnut, Manzhurskiy nut, Lancaster nut, Ailantholus nut, black nut with symptoms: drying out, wilting, ulcerative lesions of the stem.

Isolation *Phytophthor*a from soil samples was carried out by the method of floating biological baits with subsequent isolation of pathogens on a semi-selective nutrient medium-P5ARP [H].

Samples were taken from colonies similar in morphology to the genus *Phytophthora* for DNA isolation. After that, a classical PCR was set up with primers YPh1F / YPh2R and ITS4 / ITS5 and sequencing was performed.

Sequencing was carried out with samples whose product size corresponded to 450 bp. for primers YPh1F / YPh2R and 900 bp. for primers ITS4 / ITS5.

## Results

In 2020, a survey of plantings of the genus *Junglans* in the plantations of Moscow was carried out. For laboratory analysis, soil was taken from plants with signs of growth retardation and wilting, with elongated necrosis on the trunk and shoots. Isolation of late blight pathogens from soil samples was carried out by two methods: submerged sowing on a selective nutrient medium P5ARP [H] and baiting method.

After incubation of the deep-seeding plates, they were examined and the growth of colonies, characteristic of late blight pathogens, was noted. For subsequent reseeding and identification, colonies of light white color, well visible, with uneven edges, with characteristic concentric rings, were selected. Observation of the colonies during incubation showed that they differed among themselves in terms of growth rate and the nature of mycelium. Individual colonies were subcultured onto agar with carrot pieces.

At the same time, the isolation of pathogens was carried out from soil samples by the method of floating biological baits, which were selected as walnut leaves. Plates with bio-baits were incubated at 22–24°C for 5 days. For further research, cups were selected where the bait leaves were partially necrotic (Figure 2).

**Figure 1.**
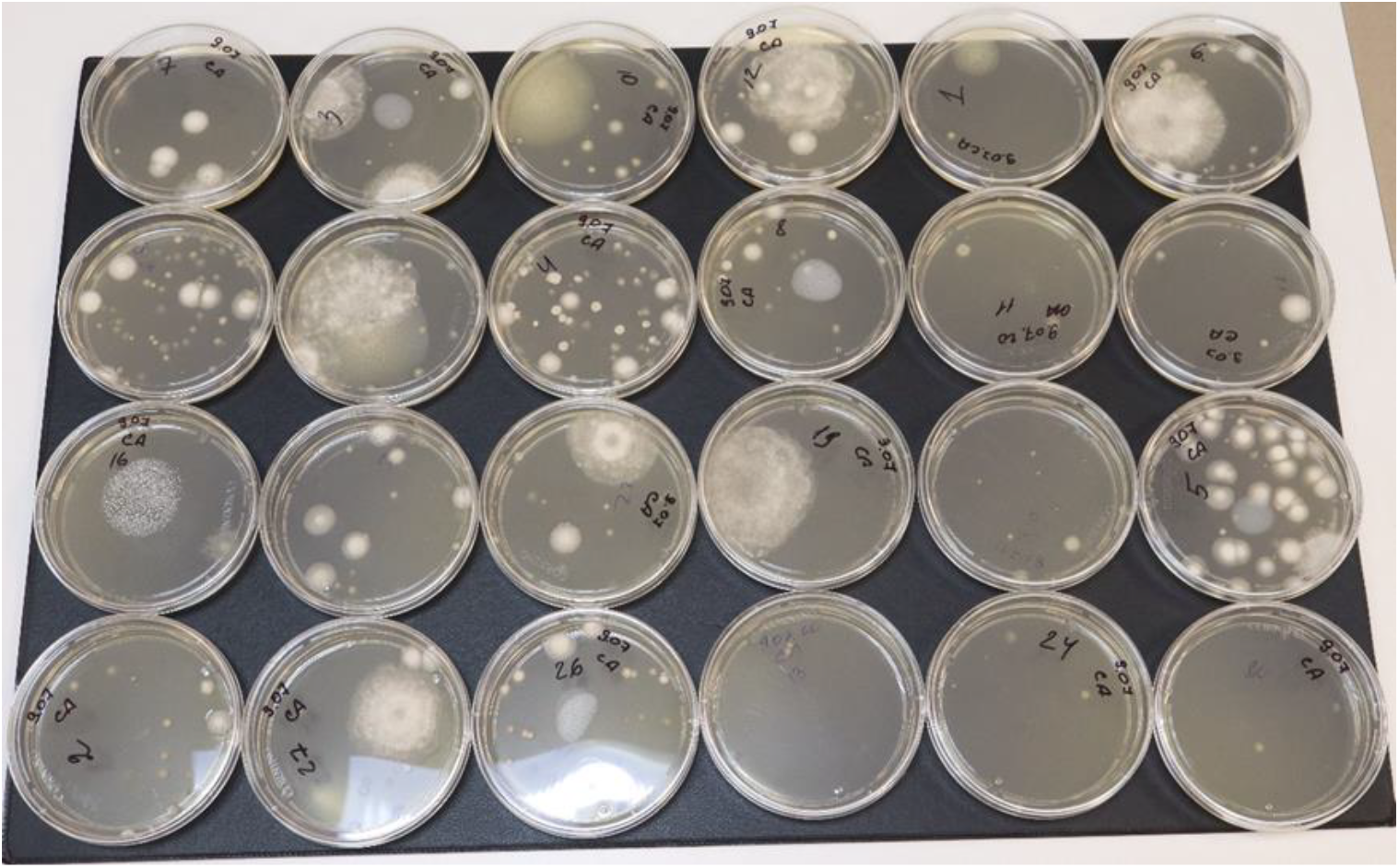
Deep sowing of soil suspension on P5ARP [H] medium (growth duration 5 days)

**Figure 2.**
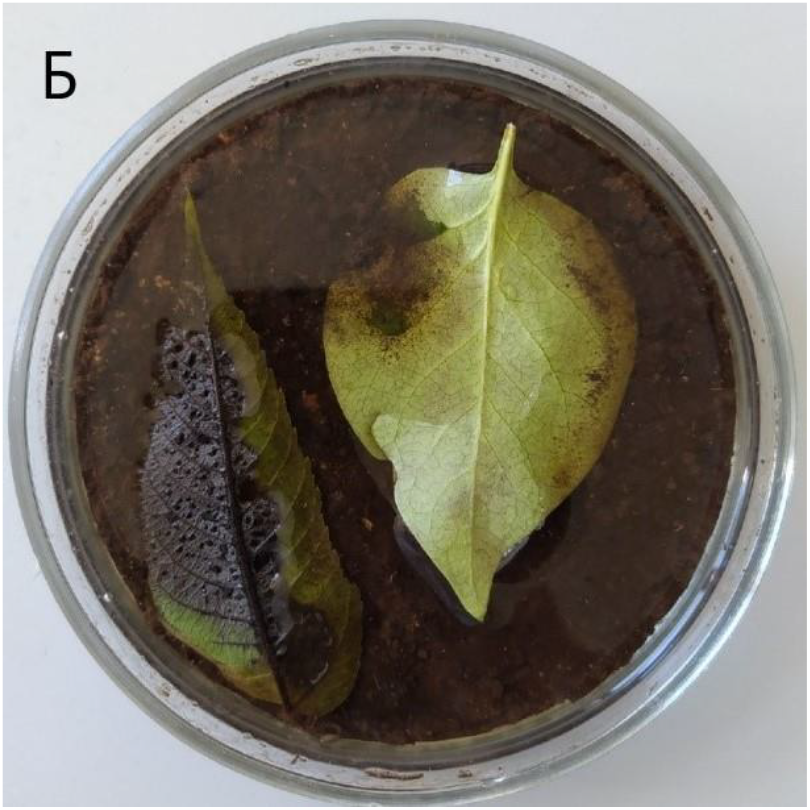
Floating bio-bait with necrosis on a walnut leaf (incubation duration 4 days)

Small fragments were excised from the diseased leaves to capture healthy and infected tissue, and a selective nutrient medium was placed on the floor. As a result, colonies with characteristic morphological features of oomycetes were formed. White, cobweb, chrysanthemum-like mycelium growth was observed on the nutrient medium (Figure 3).

**Figure 3.**
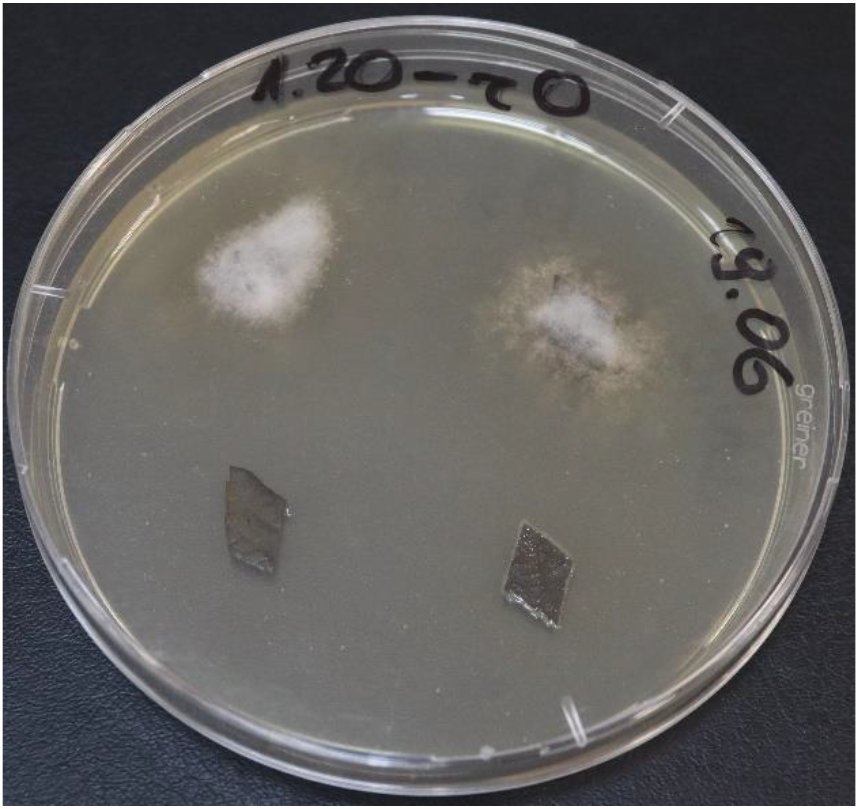
Colonies of oomycetes from bait segments on P5ARP [H] medium (growth duration 6 days)

The study of the cultural and morphological characteristics of the obtained cultures showed that the obtained isolates belong to the papillary species, presumably attributed by us to clade 1 according to H. Yang, 2017.

The identity of the isolated isolates was confirmed by deciphering the nucleotide sequence of the internal transcribed spacer (ITS) using the universal primers ITS4 and ITS5 (Cooke et al. 2000) of the Ypt1 gene (Schena et al. 2008). BLAST analysis of nucleotide sequences showed 100% identity with the published sequences of *Phytophthora cactorum* in one sample and with *P. citricola* in another with those available from GenBank.

As a result of isolation of soil oomycetes, colonies similar to the morphological characteristics of *P. cactorum* and *P citricola* were obtained.

On carrot agar, whitish colonies are clearly visible. The mycelium is appressed, creeping, white. d = 70 mm on the 7th day. Sporulation is present. The colonies are compact, slightly radiant, without a clear border, not dense with a homogeneous slightly aerial mycelium. The optimum growth temperature is 20-25 °C, the maximum temperature for growth is 31°C.

Sporangia of *P. cactorum* were oval or lemon-shaped, inversely pear-shaped, rarely asymmetrical, 20-55 × 17-46 μm in size. Oospores are spherical, 20-30 μm in diameter, colorless and yellowish with a shell 1-2 μm thick. Chlamydospores were formed in large numbers, spherical, colorless or yellowish, 19-53 microns in diameter.

The sporangia form of *P. citricola* is varying. They are obovate, obovate-clavate, pear-shaped, or slightly flattened on one side; more rarely, deeply bifurcated with two peaks or irregularly shaped. Sporangia 30-75 μm long and 21-44 μm wide (average 47 × 34 μm), non-falling and persistent on a pedicle. The semi-papillary papillae are wide and flat. Sporangiophores are simple or sometimes sympodial.

Oomycete *P. cactorum* was first isolated in 1870 from diseased cacti. This is one of the few pathogens of late blight with a wide range of affected plants - more than 250 species. The causative agent causes the death of young seedlings of ash, beech, coniferous trees; rot of fruits of apples, pears, apricots, strawberries, pumpkins; leaf and stem rot of cacti, gooseberries, rhododendrons, lilacs, ginseng, rhubarb; collar rot of apple and other fruit trees; ulcers of avocado, pear, birch, maple and oak stems; and root rot in general (Waterhouse and Waterston, 1966). *P. cactorum* is found everywhere in Europe, in addition, it is found in Asia, North America, South America, Australia, Africa, New Zealand. In the Russian Federation, the pathogen was found in the South taiga region; Forest-steppe region of the European part of the Russian Federation (Voronezh, Saratov, Kursk regions); North Caucasian mountain region (Krasnodar Krai); Priamursko-Primorsky coniferous-deciduous region (Primorsky Krai). The pathogen can cause root rot, damaging the host’s conduction system, slowing growth, causing plant death, which is especially dangerous for fruit crops. On strawberry plants, *P. cactorum* causes root rot and leathery berry rot.

*Phytophthora citricola* was first isolated from the Seccan orange fruit with brown rot symptoms in Formosa (Taiwan) and described by Sawada in 1927 (Sawada, 1927). The pathogen can also cause root rot and stem ulcers in many woody host plants.

At the beginning of this century, it was discovered that *P. citricola* is a complex of several morphologically similar species, which includes *P. plurivora, P. pini.*

*P. gonapodyides, P. cinnamomi; P. citrophthora, P. chlamydospora* species registered on walnuts and causing serious economic losses in different countries. In California, a complex of *Phytophthora* species, including *P. citricola, P. cactorum, P. cinnamomi, P. citrophthora, P. megasperma* and *P. cryptogea,* was found to cause root rot in black walnuts, resulting in tree death.

*Phytophthora cinnamomi* Rands is one of the most destructive pathogens known and can affect over 950 plant species. It is possibly the most widespread invasive organism (Hardham, 2005). *P. cinnamomi* is widespread in Europe. *P. cinnamomi* is extremely harmful. Severely affected trees are characterized by sparse, pale-colored foliage and shrinkage of individual branches. The rot surrounding the tree leads to its complete drying out. The drying process is rather slow.

